# Parkin is not required to sustain OXPHOS function in adult mammalian tissues

**DOI:** 10.1101/2023.09.02.556020

**Authors:** Roberta Filograna, Jule Gerlach, Hae-Na Choi, Giovanni Rigoni, Michela Barbaro, Mikael Oscarson, Seungmin Lee, Katarina Tiklova, Markus Ringnér, Camilla Koolmeister, Rolf Wibom, Sara Riggare, Inger Nennesmo, Thomas Perlmann, Anna Wredenberg, Anna Wedell, Elisa Motori, Per Svenningsson, Nils-Göran Larsson

## Abstract

Loss-of-function variants in the *PRKN* gene encoding the ubiquitin E3 ligase PARKIN cause autosomal recessive early-onset Parkinson’s disease (PD). Extensive *in vitro* and *in vivo* studies have reported that PARKIN is involved in multiple pathways of mitochondrial quality control, including mitochondrial degradation and biogenesis. However, these findings are surrounded by substantial controversy due to conflicting experimental data. In addition, the existing PARKIN-deficient mouse models have failed to faithfully recapitulate PD phenotypes. Therefore, we have investigated the mitochondrial role of PARKIN during ageing and in response to stress by employing a series of conditional *Parkin* knockout mice. We report that PARKIN loss does not affect oxidative phosphorylation (OXPHOS) capacity and mitochondrial DNA (mtDNA) levels in the brain, heart, and skeletal muscle of aged mice. We also demonstrate that PARKIN deficiency does not exacerbate the brain defects and the pro-inflammatory phenotype observed in mice carrying high levels of mtDNA mutations. To rule out compensatory mechanisms activated during embryonic development of *Parkin*-deficient mice, we generated a mouse model where loss of PARKIN was induced in adult dopaminergic (DA) neurons. Surprisingly, also these mice did not show motor impairment or neurodegeneration, and no major transcriptional changes were found in isolated midbrain DA neurons. Finally, we report a patient with compound heterozygous *PRKN* pathogenic variants that lacks PARKIN and has developed PD. The PARKIN deficiency did not impair OXPHOS activities or induce mitochondrial pathology in skeletal muscle from the patient. Altogether, our results argue that PARKIN is dispensable for OXPHOS function in adult mammalian tissues.

## Introduction

Pathogenic variants in the *PRKN* gene, also known as *PARK2 or PARKIN*, were identified over twenty-five years ago in patients affected by an autosomal recessive form of PD ^1,2^. Comprehensive screens of large PD patient populations have revealed that *PRKN* variants are an important cause of early-onset PD^3,4^. A variety of disease-causing mutations, including single base-pair substitutions, small and large deletions, and splice site variants, have been identified and typically lead to abolished PARKIN expression ^5^; almost 50% of variants are copy number variants ^6^. Besides homozygosity or compound heterozygosity for several loss-of-function mutations, haplo-insufficiency caused by heterozygous *PRKN* variants have been suggested to be risk factors for PD development ^7^.

PARKIN is a 465 amino acid ubiquitin E3 ligase residing in the cytosol and on the outer mitochondrial membrane (OMM), where it is involved in targeting substrates for degradation through the ubiquitin/proteasome system ^8^. Initially, it was hypothesized that loss of PARKIN would result in the accumulation of toxic substrates causing the degeneration of dopaminergic (DA) neurons and eventually PD. Although the function of PARKIN remains controversial, a wealth of data implicates PARKIN in the maintenance of mitochondrial integrity and quality control through the degradation of damaged mitochondria (mitophagy) or the regulation of mitochondrial biogenesis, or by affecting mitochondrial fusion and fission ^9^.

The consequences of PARKIN loss have been investigated by many research groups using both *in vitro* and *in vivo* models. In particular, the involvement of PARKIN in mitophagy has mainly been established by inducing ectopic high PARKIN expression in cancer cell lines lacking endogenous PARKIN expression. Treatment of these cells with a high concentration of mitochondrial uncoupling agents, e.g., CCCP/FCCP, leads to depolarization of mitochondria and other cellular compartments, including lysosomes, and to the localization of PARKIN to the OMM ^10,11^. Although this phenomenon is robustly occurring in cell culture systems, serious concerns have been raised about the *in vivo* relevance because of the ectopic expression of PARKIN and the non-physiological, near total mitochondrial membrane potential depolarization ^12,13^.

Elegant studies in *Drosophila melanogaster* have demonstrated that PARKIN plays an important role in mitochondrial maintenance in several tissues, including thoracic flight muscles, DA neurons, and sperm ^14,15^. Although PARKIN-deficient fruit flies exhibit enlarged mitochondria with disrupted mitochondrial ultrastructure^15^, experiments with pH-sensitive fluorescent mitophagy reporters have failed to unambiguously show that the mitochondrial defects are caused by impaired mitophagy ^16,17^. It thus remains unclear whether the flight muscle phenotype in *Parkin*-knockout flies is caused by defective mitophagy, accumulated mitochondrial damage and degeneration, or if it is due to impaired mitochondrial biogenesis during development.

In mice, knockout of *Parkin* does not cause DA neuron death or motor deficits ^18,19^. However, a modest increase of extracellular DA levels has been reported in the striatum ^19^ and may possibly constitute a compensatory response. Recent studies have reported that the *Parkin* knockout mice have a minor accumulation of damaged mitochondria in DA neurons ^20^ and partially impaired OXPHOS in skeletal muscle leading to myofiber atrophy ^21^. Overall, the scientific literature is confusing and the conflicting outcomes from different studies are likely a consequence of the lack of robust phenotypes in *Parkin* knockout mice. In this paper, we have performed a series of analyses in the mouse to determine how the absence of PARKIN affects OXPHOS capacity during ageing and upon mitochondrial stress. We have also investigated how *Parkin* disruption in adult DA neurons affects locomotion and transcriptional responses in this neuronal population. Our results do not support an important role for PARKIN in maintenance of the OXPHOS system in the mouse. Lastly, to investigate the relevance of these findings in human pathophysiology, we analyzed a patient with autosomal recessive PD caused by a known and a novel *PRKN* pathogenic variant that both abolish the expression of PARKIN. In this patient, PARKIN loss did not impair OXPHOS function in skeletal muscle and no signs of mitochondrial myopathy were present.

## Results

### Mitochondrial function is maintained in aged mice lacking PARKIN

There are several reports that homozygous *Parkin* knockout mice are indistinguishable from controls during their first months of life ^22–24^, but it is unclear whether the age-associated decline in mitochondrial function ^25,26^ will induce a phenotype. To study the impact of PARKIN loss in aged mice, we generated germline whole-body knockout mice by breeding heterozygous mice with a loxP-flanked exon 7 of *Parkin* (*Parkin^+/loxP^*) ^27^ to mice ubiquitously expressing cre-recombinase (β-actin-cre). This cross generated heterozygous knockouts (*Parkin^+/-^*) that were intercrossed to obtain homozygous knockouts (*Parkin^-/-^*). The β-actin-cre transgene was removed by breeding. Western blot analyses from total tissue homogenates demonstrated that the PARKIN protein was absent in brain and liver, thus confirming efficient disruption of *Parkin* (Fig. 1A). At 15 months of age, both male and female mice lacking PARKIN appeared healthy and showed no difference in body and heart weight when compared with age-matched controls (Fig. 1B).

**Fig. 1.**
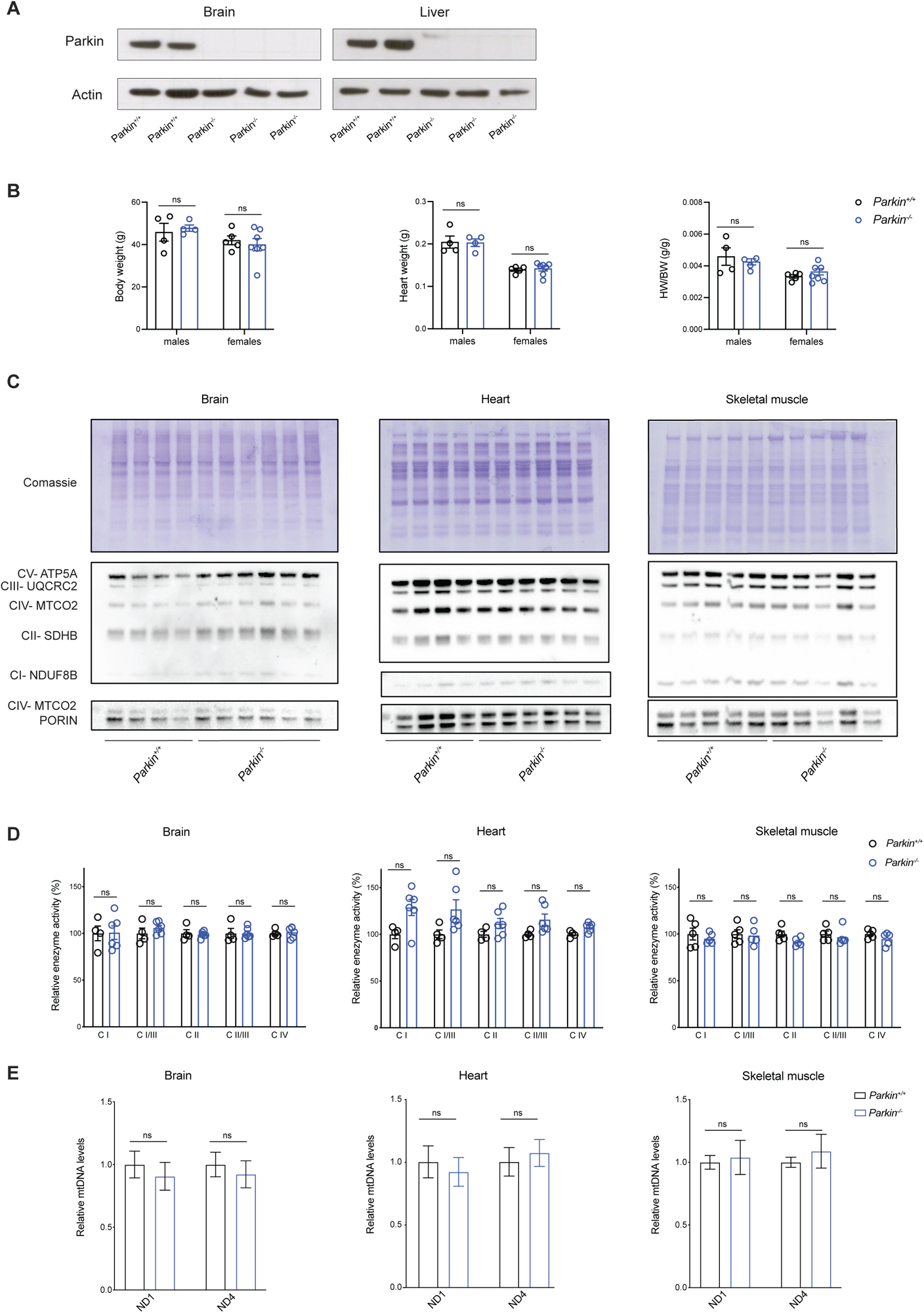
Aged *Parkin* knockout mice have normal OXPHOS function and mtDNA levels. **A.** WB analysis of PARKIN protein levels in total extracts from brain and liver of control (*Parkin^+/+^)* and KO *(Parkin^-/-^)* mice. ACTIN was used as a loading control. **B.** Body weight, heart weight and heart to body weight ratio in female and male controls and KOs at 15 months of age. Data are represented as means ± SEM; n≥ 4 per gender; ns= not significant **C.** WB analysis of OXPHOX subunits steady-state levels in mitochondrial protein extracts from brain, heart, and skeletal muscle. **D.** Respiratory chain enzyme activity assays of CI, CI + CIII, CII, CII + CIII, and CIV in isolated mitochondria from brain, heart, and skeletal muscle (gastrocnemius and soleus). Data are represented as means ± SEM; n≥ 4; ns= not significant. **E.** Quantification of mtDNA copy number performed by qPCR (ND1 and ND4/18S rRNA) in brain, heart, and skeletal muscle. Data are represented as means ± SEM; n≥ 4; ns= not significant.

We proceeded to investigate how *Parkin* ablation affects OXPHOS function in tissues with high energy demand, such as brain, heart, and skeletal muscle, of aged mice (Fig. 1C, D). No changes in steady-state levels of OXPHOS subunits were found in aged *Parkin* knockout mice in comparison with controls by using western blot analyses (Fig. 1C and Fig. S1A). Next, we assessed OXPHOS function in brain, heart and skeletal muscle and found normal respiratory chain enzyme activities in aged mice lacking PARKIN (Fig. 1D). We measured the mtDNA copy number, as an indicator of mitochondrial mass, by using qPCR analyses with two different probes (ND1 and ND4) and found no alterations in mtDNA levels of *Parkin* knockout mice (Fig. 1E), arguing against an aberrant mitochondrial turnover ongoing in the analyzed tissues. To summarize, the age-associated deterioration of mitochondrial function is not aggravated by loss of PARKIN in key energy demanding tissues of the mouse.

### *Parkin* loss does not exacerbate the phenotypes of mtDNA mutator mice

To investigate the debated role of PARKIN under mitochondrial stress conditions, we generated *Parkin* knockout mice that were also homozygous for the mtDNA mutator allele (*Parkin^-/-^*; *PolgA*^mut/mut^). The mtDNA mutator animals exhibit a premature ageing phenotype characterized by severe mitochondrial impairment caused by the progressive accumulation of mtDNA mutations ^28,29^. In two previous studies ^30,31^, loss of PARKIN was reported to cause motor impairment and degeneration of DA neurons in mtDNA mutator mice. At variance with these results, a third study ^32^ reported no aggravation of the ageing phenotypes and no loss of DA neurons when *Parkin* was disrupted in mtDNA mutator mice.

Given that maternal transmission of mtDNA mutations can be a confounding factor when mtDNA mutator mice are bred ^33^, we used a breeding strategy where heterozygous mtDNA mutator males were crossed to heterozygous *Parkin* knockout females (Fig. S1B) to circumvent this problem. This breeding scheme (Fig. S1B) generated double heterozygous mice (*Parkin^+/-^; PolgA^+/mut^*) that were intercrossed to produce four experimental groups (genotypes: *Parkin^+/+^*; *PolgA^+/+^*, *Parkin^-/-^*; *PolgA^+/+^*, *Parkin^+/+^*; *PolgA^mut/mut^*, *Parkin^-/-^*; *PolgA^mut/mut^*) that were aged for 36 weeks. At this time point, the mtDNA mutator (*Parkin^+/+^*; *PolgA^mut/mut^*) animals manifested a substantial decrease in body weight (∼20-25%) when compared with control (*Parkin^+/+^*; *PolgA^+/+^*) and *Parkin* knockout (*Parkin^-/-^*; *PolgA^+/+^*) mice. Notably, the weight loss that is typically observed in the mtDNA mutator mice ^28^ was not aggravated by *Parkin* knockout (*Parkin^-/-^*; *PolgA^mut/mut^*), arguing against an additive effect on the phenotype (Fig. 2A).

**Fig. 2.**
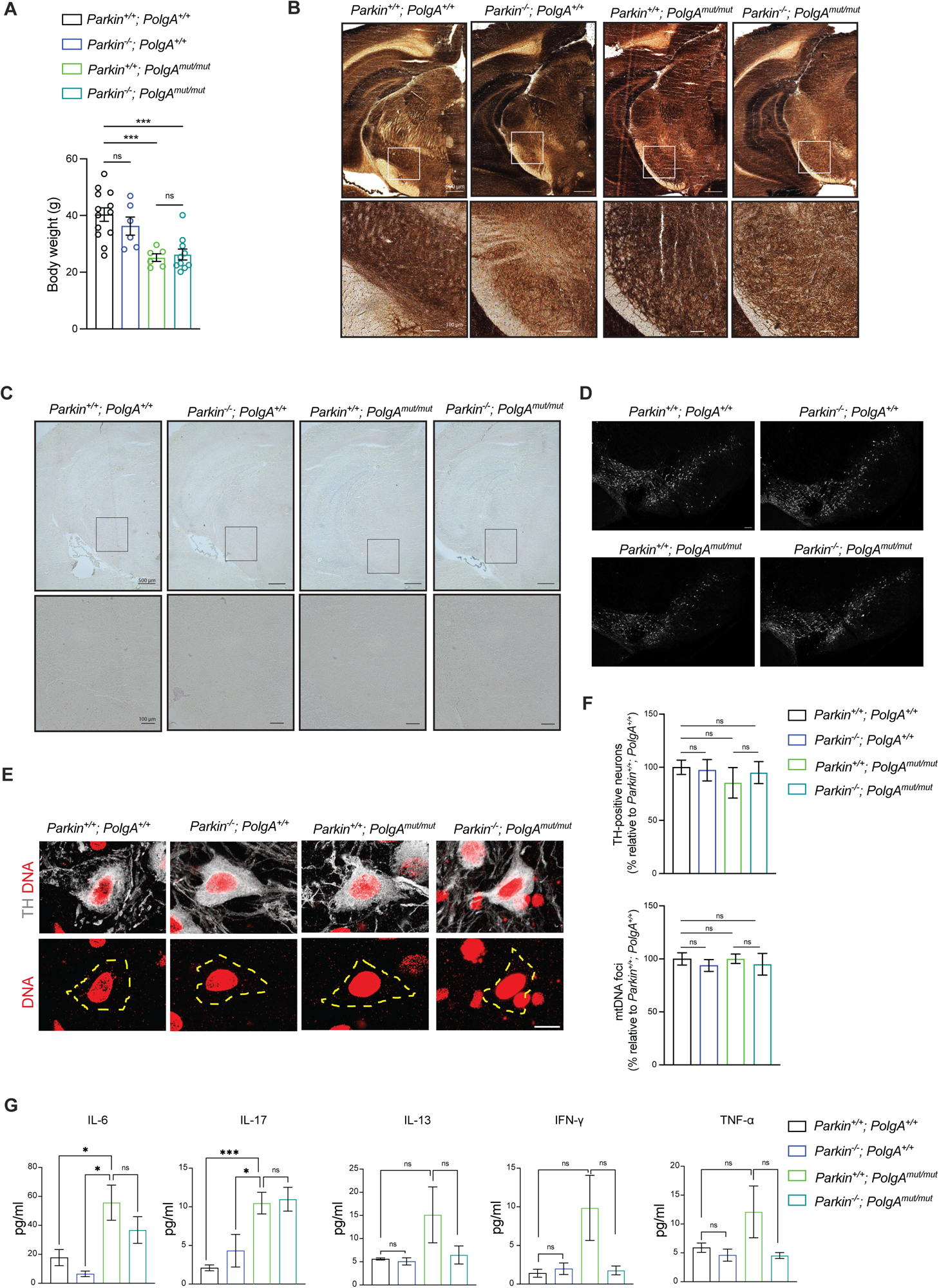
*Parkin* loss does not exacerbate the phenotypes of mtDNA mutator mouse. **A.** Body weight measured in both female and male mice at 36 weeks of age. n≥ 6 per group. **B-C.** Representative images of COX enzyme activity in ventral midbrain section in 36-week-old mice measure using **B.** COX/SDH staining (Scale bars: 500 μm and 100 μm) and **C.** NBTs histochemistry (Scale bars: 500 μm and 100 μm). **D.** Representative confocal images of TH-immunostained neuron in the midbrain sections (Scale bars: 100 μm). **E.** Representative images of DNA staining (red) in TH+ neurons (grey) of the ventral midbrain; the area occupied by TH neurons is masked in yellow (scale bar: 10 μm). **F.** Quantification of TH-positive DA neurons performed in 4-5 midbrain sections per mouse per genotype. n = 2. Quantification of mtDNA foci. n=20-42 cells from 1-3 mice per genotype. **G.** Levels of inflammatory cytokines (pg/ml) in plasma samples collected from mice at 36 weeks of age. n≥ 5 per genotype.

We proceeded to test whether *Parkin* knockout compromises OXPHOS capacity in different brain regions, i.e., hippocampus and the midbrain, of the mtDNA mutator mice. We assessed the activity of complex IV (cytochrome *c* oxidase, COX) *in situ* by employing histochemical double straining for COX and succinate dehydrogenase (COX/SDH) activity ^34^ as well as by the nitrotetrazolium blue exclusion assay (NBTx) enzyme histochemistry^35^. As a positive control, we used the hippocampal sections from mitochondrial late onset neurodegeneration (MILON) mice, which have postnatal disruption of the mitochondrial transcription factor A gene (genotype *Tfam*^loxP/loxP^; +/*CamKII-cre*) causing mtDNA depletion and OXPHOS deficiency in forebrain neurons ^36^. The MILON mice showed severe COX deficiency (blue cells on enzyme histochemistry) in the hippocampus (Fig. S2A, B) at 5 months of age, in line with our previous findings ^36^. The mtDNA mutator mice (*Parkin^+/+^*; *PolgA^mut/mut^*) presented a clear decline in COX activity in the hippocampus at 36 weeks of age (Fig. S2A, B) consistent with previous reports ^34,37^. PARKIN loss did not impact on the hippocampal COX activity in mice carrying normal mtDNA (*Parkin^-/-^*; *PolgA^+/+^*). Furthermore, the absence of PARKIN did not worsen the mitochondrial dysfunction observed in mtDNA mutator mice (*Parkin^-/-^*; *PolgA^mut/mut^*) as judged by enzyme histochemistry (Fig. S2A, B). We analyzed the midbrain and found that accumulation of mtDNA mutations (*Parkin^+/+^*; *PolgA^mut/mut^*) alone or in combination with PARKIN loss (*Parkin^-/-^*; *PolgA^mut/mut^*) did not cause any detectable decrease in COX activity *in Substantia nigra* (SN) (Fig. 2B, C). We next quantified the number of midbrain DA neurons expressing tyrosine hydroxylase (TH) in the SN and ventral tegmental area (VTA) by using confocal microscopy (Fig. 2D, F and Fig. S2C). No differences in DA neuron viability were observed in the four groups of mutant mice at age of 36 weeks. We also performed confocal analysis of brain sections to measure the number of mtDNA foci in TH positive neurons. However, we did not detect substantial changes in the number of mitochondrial nucleoids in the four groups (Fig 2E, F). Thus, loss of PARKIN in combination with increased levels of mtDNA mutations does not impact mtDNA copy number nor survival of DA neurons in the midbrain.

A recent study reported that the ablation of *Parkin* triggers inflammation induced by stimulator of interferon gene (STING) pathway in mtDNA mutator mice resulting in a profound increase in the levels of pro-inflammatory cytokines from 20 weeks of age ^31^. To examine the involvement of PARKIN in mitochondria-driven innate immunity, we measured 45 circulating plasma cytokines and chemokines in the four groups of mutant mice at the age of 36 weeks (Fig. 2G and Fig. S2C). We observed a significant increase in the levels of IL-6 and IL-17 in mtDNA mutator mice (Fig. 2G and Fig S2C). There was also a clear trend, although not significant, towards increased levels in other pro-inflammatory mediators, such as IFN-γ, TNF α and IL-13, in mtDNA mutator mice (Fig. 2G). Importantly, *Parkin* loss alone or in combination with the mtDNA mutator allele did not further increase levels of the 45 measured cytokines and chemokines (Fig. 2G and Fig S2D).

Taken together, our data show that disruption of *Parkin* does not exacerbate observed defects in the brain of the mtDNA mutator mice, which is well in line with results from a recent report ^32^. In addition, we demonstrate that *Parkin* ablation does not aggravate the inflammatory phenotype caused by progressive accumulation of mtDNA mutations.

### Induction of PARKIN loss in adult DA neurons does not cause neurodegeneration

Previous studies have suggested that *Parkin* knockout mice do not develop parkinsonism due to compensatory mechanisms occurring during development ^13,38^. This hypothesis was further supported by the observation that deletion of *Parkin* in the adult ventral midbrain, induced by stereotaxic injections of cre-expressing adenovirus, leads to a decreased mitochondrial biogenesis and progressive degeneration of DA neurons ^39,40^. To use a different and more robust approach to delete *Parkin* in midbrain DA neurons of adult mice, we generated *iParkin^DA^* (genotype: *Parkin*^loxP/loxP^, +/DatCreERT2) mice allowing the disruption of *Parkin* by tamoxifen injection (Fig. S3A) at the age of 5-7 weeks, following a previously optimized protocol ^41^. To investigate the efficiency of the inducible knockout system, we used a reporter allele expressing mitochondrially targeted YFP preceded by a loxP-flanked stop cassette (mitoYFP)^42^. As predicted, tamoxifen injection in double heterozygous (+/mitoYFP; +/Dat-cre-ERT2) mice resulted in deletion of the stop cassette and activation of YFP expression in mitochondria of TH positive (TH+) neurons (Fig. S3B). We proceeded to disrupt *Parkin* in adult DA neurons of *iParkin^DA^* mice by using tamoxifen injection and to test their motor performance over time. Surprisingly, when measuring horizontal and vertical (rearing) motor activities using an open-field arena system we found that *iParkin^DA^* mice did not exhibit defects in locomotion at 20, 40 and 60 weeks after tamoxifen injection (Fig. 3A). In agreement with the absence of motor impairment, normal levels of TH positive midbrain DA neurons were detected in *iParkin^DA^* mice at 40 and 60 weeks after tamoxifen injection (Fig. 3B, C).

**Fig. 3.**
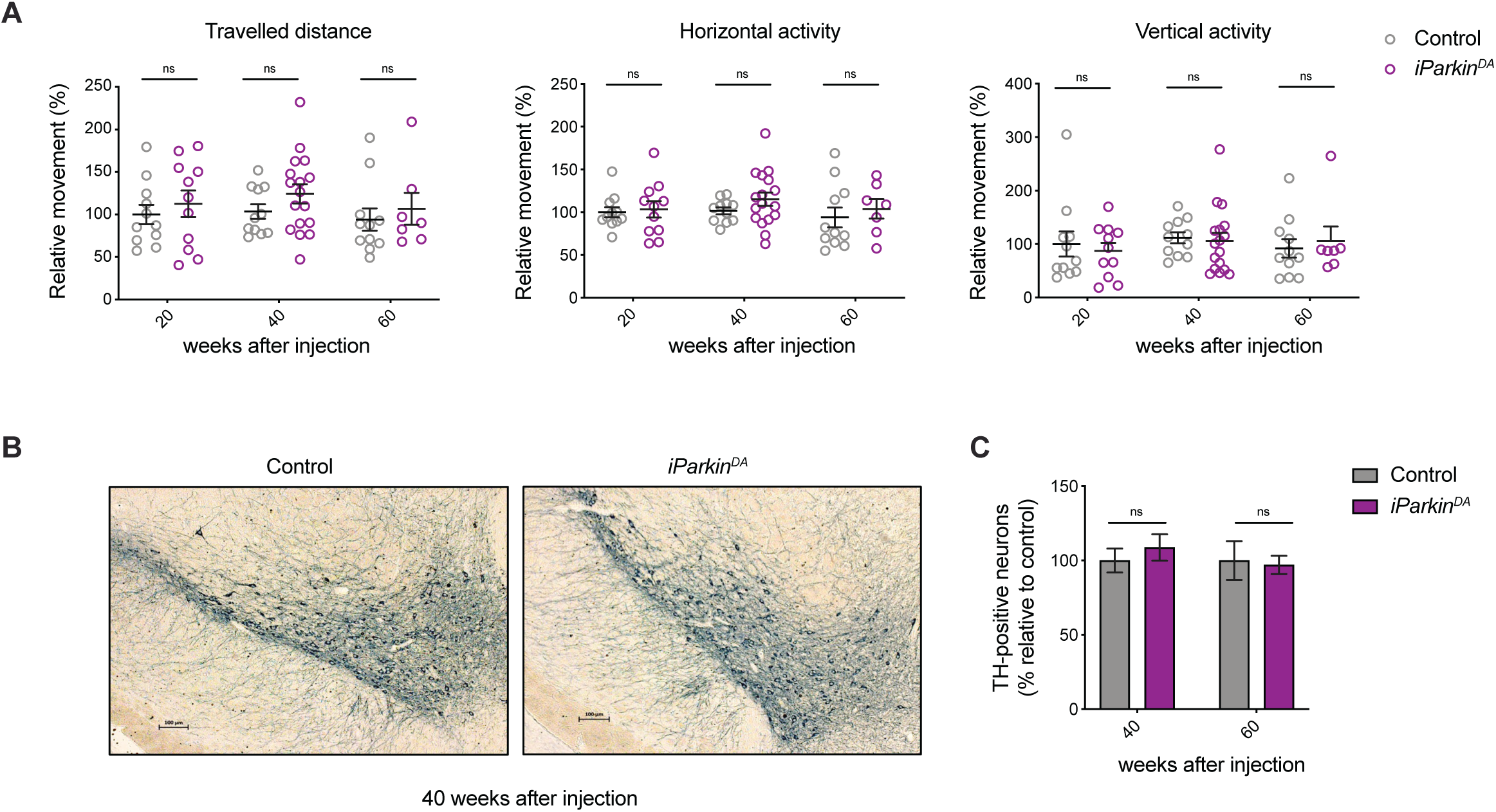
*Parkin* ablation in adult midbrain DA neurons does not lead to neurodegeneration. **A.** Spontaneous motor activity (horizontal and vertical activity and total distance) was measured in open field arena over 60 minutes. n=7-17; ns= not significant. **B.** Representative images of TH-like immunoreactivity in section from midbrain (Scale bars: 100 μm). **C.** Quantification of TH-positive DA neurons in the midbrain at 40 and 60 weeks after injection. n = 3; ns= not significant.

In summary, our results demonstrate that genetic ablation of *Parkin* in DA neurons of the adult nigrostriatal system does not result in any detectable motor defects or neurodegeneration, which argues against the hypothesis that developmental compensatory pathways are acting in *Parkin* knockout mice.

### Parkin-deficient neurons do not exhibit changes in the transcriptomic profile

It has been previously described that PARKIN regulates the levels of zinc finger protein 746 (ZNF746), also known as PARIS, via ubiquitination and degradation mediated by the proteasome system^40^. According to the proposed model, PARIS acts as transcriptional repressor controlling the expression of peroxisome proliferator-activated receptor gamma, coactivator 1α (PGC-1α), which, in turn, orchestrates important metabolic functions, including mitochondrial biogenesis, by sophisticated transcriptional mechanisms ^39^.

To investigate transcriptional responses induced by PARKIN loss in adult DA neurons we used a previously described strategy to isolate fluorophore-labelled midbrain DA neurons from adult mice ^41^. We generated *iParkin^DA^* and control mice also containing the mitoYFP allele and isolated DA neurons for bulk RNAseq analysis at 5 and 40 weeks after tamoxifen injection (Fig. 4A). For each time point, we collected mitoYFP positive and mitoYFP negative cells and prepared libraries for RNAseq analysis using Smart-seq2 protocol ^43^. Extensive quality control analyses were carried out and substantial enrichment of midbrain DA neurons was verified by the expression of well-established markers for DA neurons (e.g., *Th, Ddc*, *Slc6a3*, *Nr4a2* and *En1*) in mitoYFP positive cells from controls and knockouts, whereas specific markers expressed by astrocytes and microglia (e.g., *Gfap, Slc1a3, Aldh1l1, Itgam, P2ry12 and Tmem119*) were almost undetectable (Fig 4B, C and Fig. S3C, D). We next performed hierarchical clustering analyses using the most variably expressed genes and found that DA neurons isolated at 5 and 40 weeks after injection could not be distinguished based on PARKIN expression (Fig. 4D). Thus, the transcriptome of *Parkin*-deficient DA neurons did not exhibit major differences in comparison with controls. Additional DESeq2 analysis of the transcriptome profiles of FACS-sorted *iParkin^DA^* and control DA neurons showed no differentially expressed genes at an adjusted p value (padj) of <0.05 at 5 and 40 weeks after tamoxifen injection (Fig 4E), consistent with the observed lack of clustering (Fig. 4D). Although no significant changes were observed in the overall expression profiles of DA neurons lacking PARKIN, we decided to further interrogate our datasets by examining levels of transcripts for specific genes previously reported to be affected by PARKIN loss. This further investigation confirmed that adult-onset ablation of *Parkin* in DA neurons did not alter the expression of *Zfn746* (PARIS), *Ppargc1a* (PGC-1α) or PGC-1α-target genes, including *Nrf1* (Fig.4F and Fig. S3E). In addition, the expression levels of genes encoding inflammasome (*Nlrp3)* and Cgas (*Mb21d1*)-Sting (*Tmem173*) components were not affected by PARKIN loss at 5 and 40 weeks after tamoxifen injection. These results demonstrate that the genetic ablation of *Parkin* in adult DA neurons is not sufficient to trigger a substantial transcriptomic change that will drive or promote DA cell degeneration and death.

**Fig. 4.**
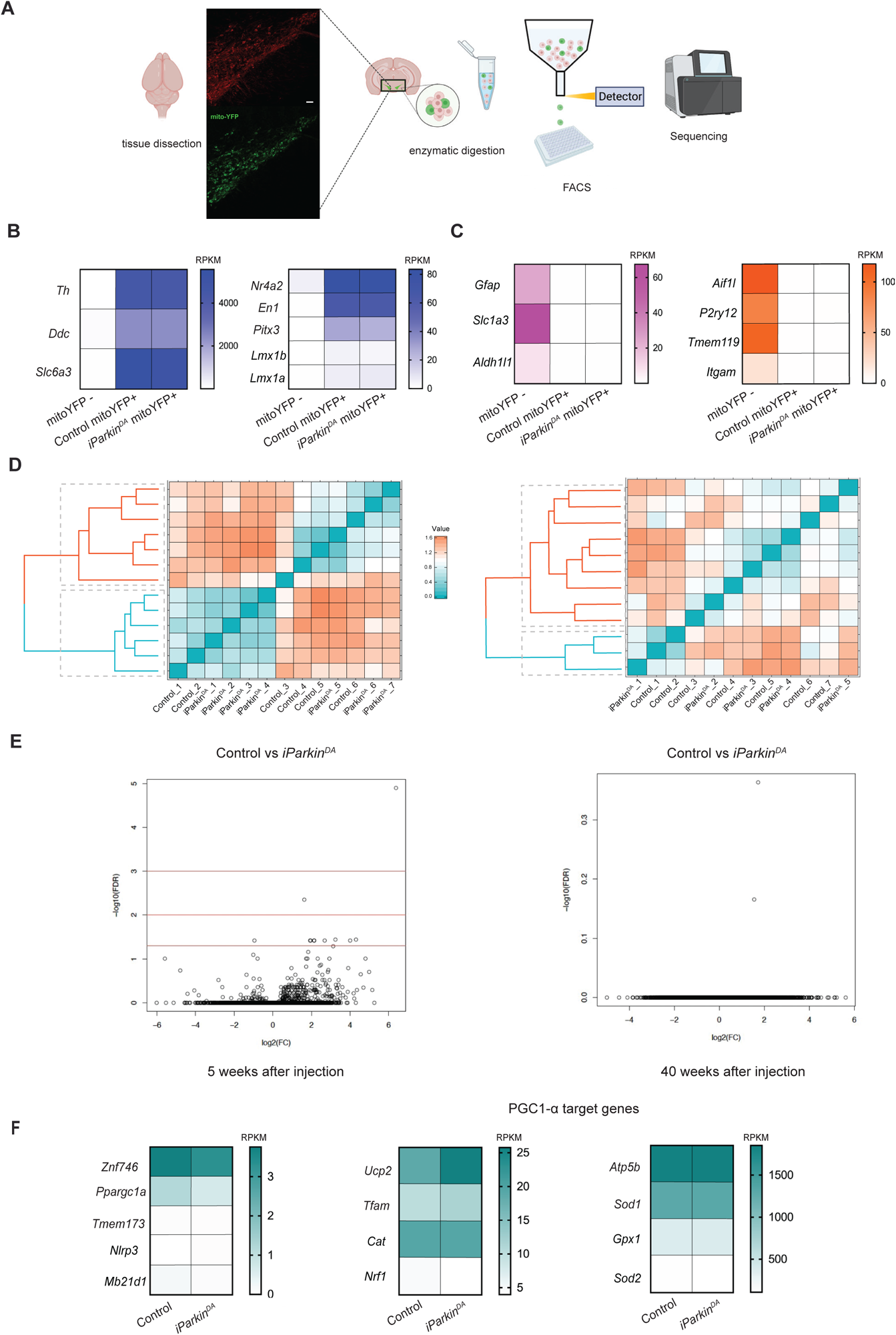
*Parkin*-deficient midbrain DA neurons do not exhibit changes in the transcriptomic profile. **A.** Experimental workflow of RNAseq in FACS-sorted DA neurons. The isolation was performed using mice expressing the mito-YFP allele under the control of the DA transporter (DAT, Slc6a3) promoter. Fluorescently labelled mitochondria (green) were visible in TH-expressing neurons (red) of the midbrain. Vibratome sections from mouse brains were enzymatically digested and mechanically triturated resulting in a single cell suspension; fluorescently-labelled midbrain DA neurons were collected by FACS; mitoYFP positive cells and mitoYFP negative were used for bulk-RNAseq. **B-C**. Heatmaps showing the expression levels of genes encoding **B**. DA neuronal markers, **C**. astrocyte and microglial markers in mitoYFP+ and mitoYFP-samples (Reads Per Kilobase Million, RPKM) at 40 weeks after injection. **D.** Hierarchical clustering analysis of RNAseq data from DA neurons isolated from *iParkin^DA^* and control mice at 5 weeks (left panel) and 40 weeks (right panel) after tamoxifen injections. n>5 per genotype and time point. **E.** Volcano plot displaying differential gene expression at 5 and 40 weeks after injections. **F.** Heatmap showing the expression of genes involved in mitochondrial biogenesis and inflammation in mitoYFP+ cells isolated from *iParkin^DA^* and control mice at 40 weeks after injection.

### Normal mitochondrial function in skeletal muscle of PARKIN deficient patient

The patient of this study presented as a teenager with slowness and stiffness. At the age of 21, she was diagnosed with suspected generalized dystonia. Her symptoms slowly worsened and at age 32, she was diagnosed with PD and initiated dopaminergic medication. She has no history of hyposmia, REM sleep behavioral disorder, depression, constipation or dysautonomia. Upon examination at the time of genetic testing at the age of 51 years, she used a walker and presented with gait difficulties with freezing episodes, slightly impaired postural control, unsteadiness, discrete bilateral rigidity, and fine movements dysfunctions. She had intermittent dyskinesias but no tremor. She had no moderate or severe non-motor symptoms and received 0 points in the non-motor symptoms scale. Her cognition was intact with a MoCa score of 27 points. Her total MDS-UPDRS score was 55 points (2+12+37+4) and Hoehn and Yahr staging 3. A brain MRI was normal, while a [123I] ioflupane SPECT examination showed major loss of the dopamine transporter (DAT) in the basal ganglia (Fig. 5A). Automated quantification using the BRASS program showed reduced DAT uptake in nucleus caudatus and putamen bilaterally. Her symptoms were responsive to a combination of low doses of levodopa, COMT inhibitor and D2R dopamine agonist.

**Fig. 5.**
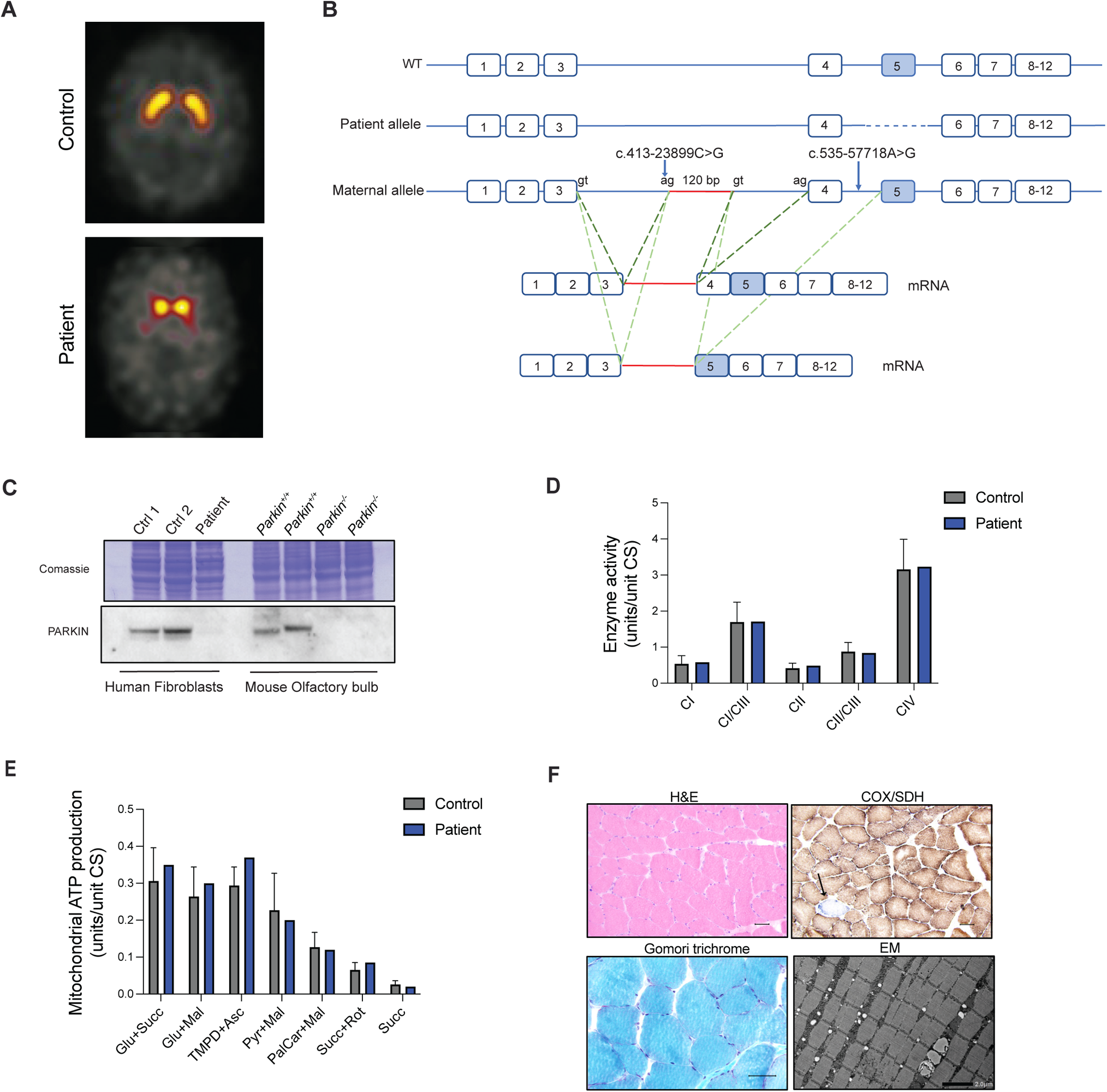
Loss of PARKIN in a PD-patient does not affect the OXPHOS function in skeletal muscle. **A** DaTSCAN® SPECT of presynaptic dopaminergic terminals in the striatum in control and the PARKIN deficient PD patient. **B.** Wild type *PRKN* gene and alleles identified in the patient. Results of the mRNA analysis of the maternal allele are shown. In red the 120bp intronic sequence inserted in the mRNA. Additional AG and GT splicing sites are indicated. **C.** WB analysis of PARKIN steady-state levels in total protein extracts from control subjects and *PARKIN* PD patient. Total extracts from olfactory bulb of control (*Parkin^+/+^)* and KO *(Parkin^-/-^)* mice were used as reference. **D.** Enzyme activity assays of CI, CI + CIII, CII, CII + CIII, and CIV in isolated mitochondria from skeletal muscle from controls and the patient. **E.** The mitochondrial ATP production rate in isolated mitochondria from skeletal muscle from controls and the patient**. F.** Skeletal muscle histopathology using staining for COX/SDH, hematoxylin and eosin (H&E), Gomori trichrome. Transmission electron microscopy analysis was performed to define the distribution and morphology of mitochondria in skeletal muscle from the patient.

A detailed genetic analysis on patient biopsy performed by whole genome sequencing identified a heterozygous 11,6 kb deletion that includes the entire exon 5 of *PRKN* with an insertion of 34 nucleotides at the breakpoint; this variant is described as c.575 4062_658 + 7471delinsTAAATTAGAAAATTAATTAGAAAAATTTAAATTA, p.(Gly179_Ala206del). No other pathogenic variants were identified in the coding region or at the exon/intron boundaries. Further investigations evaluating deep intronic changes revealed the presence of two uncommon variants that were predicted to create a possible splice site: c.413-23899C>G (g. 162646183C>G) and c.535-57718A>G (g. 162532924A>G) in intron 3 and intron 4, respectively. These intronic variants were maternally inherited, while the exon 5 deletion was not present in the mother but unfortunately the DNA from the father was not available. To understand whether these variants affect *PKRN* expression, we performed RNA analysis that showed that the patient does not express a wild-type mRNA. Using primers targeting exon 1 and 7, three different RT-PCR products were amplified: one missing exon 5, as a consequence of the exon 5 deletion presumably inherited from the paternal allele, and two PCR products presented with a 102 bp sequence insertion starting from the nucleotide after the variant c.413-23899G>C. Notably, one of these two amplicons also lacked exon 4 (Fig 5B). The same analysis performed on the maternal sample showed the presence of a wild-type product as well the two aberrant splicing products.

Although it remains unclear whether the exon skipping is due to variant c.413-23899G>C or variant c.535-57718A>G in intron 4, we proceed to investigate how the presence of the described compound heterozygous variants impacts on PARKIN protein steady-state levels. A western blot analysis of total protein extracts from fibroblasts of the patient showed absence of the PARKIN protein (Fig. 5C), confirming that the deep intronic variants indeed generate a loss-of-function allele. To study the effect of PARKIN loss in adult human tissues and mitochondria, a muscle biopsy was obtained from the anterior tibial muscle of the patient using a conchotome and mitochondria were isolated. Assessment of the OXPHOS function showed normal respiratory chain enzyme activities (Fig. 5D) and normal ATP production rates (Fig. 5E) in skeletal muscle. In addition, histopathological analyses revealed that skeletal muscle tissue was well preserved without any necrotic fibers or inflammatory cell infiltrates (Fig. 5F). COX/SDH staining identified the presence of a single COX-deficient fiber (Fig. 5F), compatible with the age of the patient and in line with normal NADH and SDH staining (Fig. S4). Staining for glycogen and neutral lipid were also normal (Fig. S4) and no ragged-red muscle fibers were detected using Gomori trichrome staining (Fig. 5F). In addition, transmission electron microscopy showed no alterations in muscle morphology and mitochondrial ultrastructure (Fig. 5F and Fig. S4).

## Discussion

Although our results argue against a critical function for PARKIN in maintaining OXPHOS capacity, it should be emphasized that PARKIN is highly conserved across species ^44^ and is present in both vertebrates, e.g., human, rat, mouse, bird, frog, and invertebrates, e.g., fruit-flies. The gene encoding PARKIN has thus been maintained by selection during evolution of metazoa and must therefore have an important function. Biochemically, PARKIN is a ubiquitin E3 ligase that meditates the covalent binding of ubiquitin or ubiquitin chains to lysine residues of proteins to mark them for degradation by the proteasome or lysosome. PARKIN consists of an N-terminal ubiquitin-like (Ubl) domain and four zinc-finger domains, RING0, RING1, IBR and RING2. Several crystal structures of PARKIN and its different domains have been solved with various degrees of resolution ^45–48^, which have clarified important molecular aspects of PARKIN activation and regulation. In addition, these structural studies have significantly contributed to unveil how loss-of-function mutations can disrupt protein function by compromising the zinc coordination, the protein folding, the catalytic site, or the protein contact sites ^49^. Affinity purification techniques coupled with quantitative mass spectrometry analyses have shown that PARKIN can interact and promote the ubiquitination of a multitude of substrates without targeting a specific consensus sequence or a structural recognition motif, suggesting that the proximity to the substrate is important for the ubiquitination by PARKIN ^50^. This hypothesis has been further supported by observations that even non-mitochondrial proteins targeted to mitochondria are ubiquitinated by PARKIN ^51^.

The broad range of PARKIN substrates has linked its function with the regulation of mitochondrial integrity. In fact, PARKIN can ubiquitinate both mitofusin 1 and 2 (MFN1 and MFN2), which are the main the players in mitochondrial fusion, and target them for proteasomal degradation ^50,52–54^. In addition, PARKIN has been implicated in the regulation of mitochondrial biogenesis, primarily through the ubiquitination and subsequent proteasomal degradation of PARIS, which is a PGC1-α repressor ^39,40^. Furthermore, PARKIN, together with the serine-threonine kinase PINK1, has been reported to function as a crucial regulator of mitochondrial degradation via mitophagy. According to this model, PARKIN is activated by PINK1 in response to mitochondrial damage and thereafter ubiquitinates a variety of OMM proteins. This process is supposed to mark functionally impaired mitochondria for engulfment by the autophagosome followed by degradation in the lysosome ^55^. However, *in vivo* studies in fruit flies and mice have questioned the contribution of PINK1/PARKIN-mediated mitophagy under basal conditions ^16,56^ or in response to mitochondrial damage ^42^. Interestingly, a recent study conducted using human iPSC-derived DA neurons proposes a novel function of PARKIN in promoting mitochondrial and lysosomal amino acid homeostasis through stabilization of inter-organelle contacts ^57^.

In this work, we addressed fundamental questions about the role of PARKIN in mitochondria during ageing or upon mitochondrial stress. We demonstrate here that loss of *Parkin* in the mouse does not aggravate the naturally occurring decline in mitochondrial function of high-energy-demanding tissues. Our findings argue that PARKIN is not required to sustain tissue bioenergetics during ageing. In addition, *Parkin*-deficient mice present normal mtDNA levels, suggesting that PARKIN loss does not cause drastic changes in mitochondrial mass. Hence, these results do not support a major involvement of PARKIN in mitochondrial quality control, which suggests the existence of PINK1/PARKIN-independent pathways acting in the regulation of the fine balance between mitochondrial degradation and biogenesis. Recent reports have shown that loss of FBXL4, which is a subunit of the Skp1-cullin1-F-box (SCF) E3 ubiquitin ligase protein complex, increases mitophagy ^58^. Loss of FBXL4 decreases the mitochondrial mass with concomitant mtDNA depletion and mitochondrial dysfunction in affected tissues in humans ^59–61^ and mice ^58^. Three independent recent reports have shown that FBXL4 actively suppresses mitophagy by degrading the mitophagy receptors NIX and BNIP3 ^62–64^. The absence of FBXL4 thus increases the levels of these receptors leading to increased mitophagy, which, in turn, causes mtDNA depletion and impaired OXPHOS function in affected patients. In contrast to PARKIN, FBXL4 and its downstream targets thus provide an important example of a clinically relevant mitophagy pathway whose impairment causes severe mitochondrial disease.

It has been reported that stress induced by mtDNA mutations results in activation of the STING-mediated innate immune response when PARKIN is absent, leading to degeneration of DA neurons and motor impairment in mice ^30,31^. Using a variety of molecular and biochemical analyses, we show here that PARKIN loss does not exacerbate the proinflammatory response or neurodegeneration of mtDNA mutator mice consistent with the results from a recent independent study ^32^. Importantly, also in *Drosophila* the role of STING in the induction of *Parkin* mutant phenotypes remains controversial. In fact, two recent investigations have generated conflicting results on the impact of STING loss in suppressing behavioral and mitochondrial defects manifested by *Parkin* deficient fruit flies ^65,66^.

Homozygous *Parkin* knockout mice do not develop parkinsonism and this failure to reproduce the human phenotype has been attributed to compensatory mechanisms activated during development. In line with this proposal, it has been reported that PARKIN loss induced by stereotaxic injection of adenovirus expressing cre-recombinase in adult mice homozygous for a lox P-flanked *Parkin* allele results in mitochondrial dysfunction and DA neuron death triggered by the transcriptional repression of PGC-1α ^39,40^. We generated mice lacking PARKIN in adult midbrain DA neurons by using a tamoxifen-inducible cre-recombinase expressed under control of the DA transporter (DAT, *Slc6a3*) promoter. Surprisingly, these inducible knockout mice preserve their motor abilities and have normal levels of TH+ neurons in the midbrain. Moreover, RNA-seq analyses of isolated *Parkin*-deficient DA neurons did not reveal substantial transcriptional changes, including the transcript levels of PGC-1α-target genes. It is unclear why our results are at variance with the previous reports showing a clear phenotype after adult-onset knockout of *Parkin*. A likely explanation of the discrepancy between our results and the previous reports may lie in the different modes of expressing cre-recombinase (virus vs transgene). In fact, while the transgenic approach is genetically specific, viral approaches may cause recombination in other cell types besides DA neurons or have toxic effects when injected at higher titer doses.

In summary, we here provide compelling evidence that PARKIN is dispensable for the maintenance of OXPHOS function in adult mouse tissues, including DA neurons. These findings have been further validated in a PARKIN-deficient patient. By using use whole genome sequencing, we found that the analyzed patient carries a large deletion encompassing exon 5 and two deep intronic variants, which leads to PARKIN loss. While other exon 5 deletions were previously described in PD patients^67,68^, to our knowledge the pathogenicity of deep intronic variants in the *PRKN* gene was never demonstrated before. The only described splicing variants (see HGMD® Professional 2023.2) are either located at the canonical splice site or within 20 nucleotides from the intron/exon junction. Therefore, this case highlights the importance of the use of whole genome sequencing and rare intronic variants analysis to improve the diagnosis of *PKRN*-PD patients. Importantly, the patient of this study present normal tissue morphology and preserved respiratory chain function in skeletal muscle three decades after onset of PD symptoms, which support that PARKIN is not required to maintain OXPHOS function in adult human skeletal muscle and corroborate our findings in the mouse.

**Fig. S1. Ablation of *Parkin* in mouse models. A.** Densitometric quantification of the steady-state levels of OXPHOS subunits in brain, heart, and skeletal muscle as determined by WBs. Data are represented as mean ± SEM; n≥4. **B.** Breeding strategy used to generate the transgenic mouse line homozygous for both the whole-body *Parkin* knockout allele and the mtDNA mutator allele.

**Fig. S2. Molecular phenotypes of *Parkin^-/-^; PolgA ^mut/mut^* mice. A-B.** COX enzyme activity measured using **A.** COX/SDH and **B.** NBTx staining in brain sections of MILON mice (conditional *Tfam KO* in forebrain neurons, *cTFAM^-/-^*) at 5 months of age and in *Parkin* KO mice carrying WT mtDNA (*Parkin^-/-^; PolgA^+/+^)* or *mutant mtDNA (Parkin^-/-^; PolgA^mut/mut^)*. **C.** Quantification of TH-positive DA neurons in SN and VTA performed in 4-5 midbrain sections per mouse per genotype. n = 2. **D.** Heatmaps showing the levels (pg/ml) of 45 inflammatory cytokines and chemokines in plasma samples collected from 36-week-old mice. n≥ 5 per genotype.

**Fig. S3. Induction of PARKIN loss in adult DA neurons and transcriptomics analysis at 5 weeks after tamoxifen injection. A.** Diagram depicting tamoxifen-induced inactivation of *Parkin* gene in adult mice. Mice at 5–7 weeks of age were intraperitoneally injected with tamoxifen for 5 consecutive days and examined up to 60 weeks after injection**. B**. Representative confocal microscopy images of mitoYFP-labelled mitochondria (green) in TH immunoreactive neurons (red) at 5 weeks after tamoxifen injection (Scale bar: 20 μm)**. C-D.** Heatmaps showing the expression levels of genes encoding **C**. DA neuronal markers, **D**. astrocyte and microglial markers in mitoYFP+ and mitoYFP-samples (Reads Per Kilobase Million, RPKM) at 5 weeks after injection. **E.** Heatmap showing the expression of genes involved in mitochondrial biogenesis and inflammation in mitoYFP+ cells isolated from *iParkin^DA^* and control mice at 5 weeks after injection. n>5 per genotype.

**Fig. S4. Histochemistry and histology of** skeletal muscle from the **PARKIN-deficient patient.** A skeletal muscle biopsy specimen from *PKRN* patient stained with myosin to discriminate between type 1 fibers (red) and type 2 fibers (white), Periodic acid schiff (PAS) staining, to detect glyocogen. Staining for succinate dehydrogenase (SDH) and nicotinamide adenine dinucleotide (NADH, to reveal myofibrillar architecture). (Scale bar: 50 μm). High magnification transmission electron microscopy analysis was performed to define mitochondrial ultrastructure in skeletal muscle (Scale bar: 500 nm).

## Materials and Methods

### Ethics statement

All animal procedures were conducted in accordance with European, national, and institutional guidelines and protocols were approved by the Stockholms djurfo rso ksetiska na mnd, Sweden and by the Landesamt fu r Natur, Umwelt und Verbraucherschutz Nordrhein–Westfalen, Germany. Animal work was performed in accordance with the recommendation and the guidelines of the Federation of European Laboratory Animal Science Associations (FELASA).

The clinical part of the study was approved by the regional ethics committee in Stockholm and the Swedish ethical review authority (2019-04967). Informed consent was obtained from the patient.

### Mouse models

Homozygous mice for a *LoxP*-flanked Parkin allele (*Parkin^loxP/lo^*^xP^) were crossed to heterozygous β-actin-cre or DATcreERT2 mice in order to generated germline whole-body knockout (*Parkin^-/-^*) and adult conditional knockout (*iParkin^DA^*) mice, respectively. At 5-7 weeks of age, *iParkin^DA^* and control mice were treated for 5 consecutive days by intraperitoneal injection of 2 mg of tamoxifen (Sigma T5648 dissolved in ethanol and sunflower oil) to induce the ablation of *Parkin* gene in adult DA neurons. *PolgA* mutator allele (*PolgA^mut/mut^*), expressing the exonuclease-deficient PolgA, ^28^ and the stop–mito–YFP allele ^42^, expressing a mitochondrially targeted YFP, were subsequently introduced via additional crossing. Analyses of controls and KO mice were performed at different time points. All mice were on the C57BL/6N genetic background.

### Western Blot

Isolation of mitochondria from mouse heart, skeletal muscle was performed by differential centrifugation ^69^. Total extracts from control and patient fibroblasts were prepared in RIPA buffer supplemented with protease inhibitors (Complete, Roche). Five micrograms of isolated mitochondria or twenty micrograms of total protein extracts were resuspended in Laemmli buffer, run on 12% SDS–polyacrylamide gel electrophoresis (Invitrogen) and then transferred onto nitrocellulose membrane using *iBlot2* system (Invitrogen). Immunodetection was performed according to standard techniques using enhanced chemiluminescence Immun-Star HRP Luminol/Enhancer (Bio-Rad). The following antibodies were used: OXPHOS Rodent Antibody Cocktail (ab110413, Abcam) for NDUFB8 (CI), SDHB (CII), UQCRC2 (CIII), MTCO1 (CIV) ATP5A (CV), Porin (ab14734, Abcam), Parkin (ab15954, Abcam). Quantification of WB signals was performed by densitometry using software FIJI (ImageJ2; Version 2.3.0/1.53f).

### Biochemical evaluation of respiratory chain function and ATP production

Respiratory chain enzyme activities and ATP production were measured in isolated mitochondria from different mouse and human tissues, as previously described ^70^.

### mtDNA copy number

Genomic DNA was isolated from snap-frozen tissues using the DNeasy Blood and Tissue Kit (Qiagen), according to the manufacturer’s instructions. Quantification of mtDNA copy number was performed in triplicates using five ng of DNA using TaqMan Universal Master Mix II and TaqMan probes from Life Technologies. The mtDNA levels were assessed using probes against the mitochondrial genes (ND1 and ND4), and nuclear 18*S* rRNA gene was used as a loading control.

### Motor performance

Motor activity of control and *iParkin^DA^* mice was measured in open field arenas (VersaMax, AccuScan Instruments) at 20, 40, and 60 weeks after tamoxifen injection. Following an acclimation period of at least 30 minutes in the ventilated experimental room, mice were placed individually in activity cages (40 × 40 cm and 30 cm high) for 60 minutes. A grid of infrared light beams was used to record spontaneous horizontal and vertical activities and the total distance traveled was calculated.

### Cytokine and chemokines quantification

After cervical dislocation, blood was collected by cardiac puncture in anticoagulant-treated tubes (EDTA tubes BD) and plasma (supernatant) was obtained after centrifugation at 2000 g for 10 minutes at 4 °C. Plasma aliquots were snap-frozen in liquid nitrogen and stored at -80°C until the analysis. Mouse Cytokine/Chemokine were quantified using the 45-Plex Discovery Assay® Array (MD45) by Eve Technologies.

### Immunohistochemistry and confocal microscopy

Anesthetized mice were perfused with by 4% paraformaldehyde in phosphate buffer. The brains were dissected, postfixed for 2 hours, and equilibrated with 10% sucrose. Brains were frozen and cryo-sectioned to obtain 14 to 20 μm thick sections. After 1 hour in blocking solution (PBS+ 0.3% Triton X-100+ 1% BSA), the tissue sections were immunolabeled overnight with primary antibodies against TH (1:1000, Chemicon). For fluorescent staining, Cy3-(1:400, Jackson Biolabs), Alexa 546-(1:400, Life Technologies) and Alexa 633-conjugated secondary antibodies (1:400, Life Technologies) were used. Confocal images were acquired by sequential scanning using a LSM800 or LSM880 microscope (Zeiss).

Alternatively, fixed brains were transferred in PBS and sectioned into 50-um thick coronal slices on a Leica vibratome (Leica Microsystems GmbH, Vienna, Austria). Free-floating slices were placed in tris-EDTA buffer (pH 9) for 30 minutes at 80°C for antigen retrieval and were then blocked for 15 minutes with 3% BSA in PBS at room temperature (RT). Slices were subsequently incubated overnight at 4°C under agitation with the primary antibodies: anti-tyrosine hydroxylase (1:500, Synaptic System) and anti-DNA (1:200, Progen). Sections were then incubated with secondary antibodies conjugated with Alexa 488 and Alexa 594 at a dilution of 1:500 for 2 hours at RT. Afterwards, slices were thoroughly washed in PBS, counterstained with DAPI, and mounted on microscopic slides with AquaPolymount. Slices were imaged with a laser scanning confocal Stellaris 5 or TCS SP8 gSTED equipped with a white light laser and a 405-diode ultraviolet laser (Leica Microscope). Image acquisition was done using the LAS-X software in accordance with the Nyquist sampling sequential mode by exciting the fluorophores and collecting their signal with hybrid detectors (HyDs).

### Image Processing

Acquired images were deconvolved using the Hyugens Professional (Huygens Pro, Scientific Volume Imaging) to improve contrast and resolution. For further image processing Image J and Adobe Photoshop were used to adjust brightness and contrast. For figure preparation Adobe Illustrator was used.

### Quantification of TH+ neurons

Every sixth midbrain cryo-section (20 μm thickness) was immunolabeled for TH. For the non-fluorescent labeling, a biotinylated secondary antibody (1:400, Vector Laboratories) was used and the signal was detected by using a peroxidase substrate (Vector SG, Vector Laboratories). Nuclei of TH-positive neurons were counted in both right and left hemisphere from 9-11 sections for brain. For experiments involving vibratome sections, dopamine neurons cell density was assessed with the Cell Counter plugin (Image J software). Briefly, density values were assessed in 4-5 midbrain sections per genotype by dividing the number of counted dopaminergic neurons (TH positive cells) by the area occupied by the counted cells.

### Dual COX/SDH enzyme histochemistry

Brains from the four experimental group were rapidly collected and frozen on dry ice. Brain sections (20 μm) were stained as previously described ^71^.

### NBTx enzyme histochemistry

NBTx enzyme histochemistry was performed as previously reported ^35^. Briefly, 20 µm brain sections were equilibrated 5 minutes in PBS. Slides were then incubated at room temperature in freshly prepared NBTx staining solution (15 mM Nitrotetrazolium Blue Chloride, 0.2 mM Phenazine Methosulfate, 130 mM Sodium Succinate in PBS pH 7). After 90 minutes, samples were washed three times in PBS, dehydrated in increasing concentration EtOH solutions (50%, 70%, 96%, 100%) and in xylene and finally mounted in Cytoseal.

### Isolation of DA neurons by FACS

Brains were dissected from mitoYFP expressing mice, sectioned and dissociated into single cell suspensions, as previously described ^72^. After dissociation, mitoYFP positive and negative cells were collected using a BD FACSAria III Cell Sorter for RNAseq analysis.

### Library preparation and sequencing

FACS sorted cells (n>5 per genotype) were used to generate the cDNA libraries according to the Smartseq2 protocol as previously described ^43^. The Nextera XT DNA library preparation kit (FC-131-1024) was used for cDNA tagmentation. The quality of cDNA and tagmented cDNA was checked on a High-Sensitivity DNA chip (Agilent Bioanalyzer). Sequencing was performed on Illumina HiSeq 2500, giving 51 bp reads after de-multiplexing.

Reads were aligned to the mouse genome (mm10) merged with eGFP and ERCC spike-in sequences using Star v2.3.0 and filtered for uniquely mapping reads. Gene expression was calculated as read counts and as reads per kilobase gene model and million mappable reads (RPKMs) for each transcript in Ensembl release 75 using rpkmforgenes. Read counts were summed across technical replicates and RPKMs were averaged across technical replicates, resulting in gene expression data. Protein-coding genes were selected for further analyses.

Hierarchical clustering was performed in R using the ward.D2 agglomerative method and the Pearson correlation-based distance measure. Differential gene expression^73^ analysis was performed using DESeq2 ^74^.

### DaTSCAN SPECT

Assessment of presynaptic dopamine transporter with ^123^I-labeled N-(3-fluoropropyl)-2β-carbomethoxy-3β-(4-iodophenyl)nortropane (FP-CIT) (i.e. DaTSCAN**®)** was done at the level of striatum according to standard clinical procedure ^75^.

### Genetic Investigation

Whole genome analysis was performed on a Illumina platform described ^76^. Splicing predictions were performed by the software Alamut Visual Plus version 1.7 © SOPHiA GENETICS. RNA was extracted by whole blood collected on PAXgene™ Blood RNA tubes (Becton Dickinson) by the PAXgene™ Blood RNA Kit (Qiagen). The High Capacity cDNA Reverse Transcription Kit (Applied Biosystem) was used to synthetize cDNA; primers for RT-PCR were designed within exon1 (PRKN-1F: 5‘-CACCTACCCAGTGACCATGA-3’) and exon7 (PRKN-7R: 5‘-CTGCCGATCATTGAGTCTTG). Sanger sequencing was performed using the Big Dye Terminator v3.1 Cycle Sequencing Kit (Applied Biosystems) and the DNA analyzer 3500XL (Applied Biosystems). The reference sequence NM_004562.3 was used for *PRKN* gene.

### Histopathological tissue analysis

Skeletal muscle biopsy from the anterior tibial muscle was snap-frozen in isopentane cooled by dry ice for cryostat sectioning. Standard techniques were applied for histological stains and enzyme histochemistry including the oxidative enzymes NADH, SDH, COX and COX-SDH ^77^. A small specimen was fixed in buffered glutaraldehyde and embedded in epoxy resins for electron microscopy according to standard procedures.

### Statistical analysis

All statistical analyses were performed using GraphPad Prism v9 software. All data are in the figures are presented as mean ± SEM. Statistical comparisons were performed using single or multiple Student’s t-test or one-way analysis of variance (ANOVA).

## Acknowledgments

The authors wish to thank Dr. Valina Dawson at Johns Hopkins University, who provided *Parkin^loxP/lo^*^xP^ mice. The authors wish to thank Fredrik Holmstro m and Linda Gillberg, (from Perlmann’s lab, Karolinska Institutet) for the excellent technical assistance with tissue dissociation and FACS sorting. This study was supported by grants to NGL from Vetenskapsra det (2015-00418), Knut och Alice Wallenbergs Stiftelse, Hja rnfonden and Parkinsonfonden. RF was supported by grants from Vetenskapsra det (2022-01477), Loo och Hans Ostermans stiftelse, A hle n-stiftelsen, KI Research Foundation Grants, StratNeuro and Hedlunds stiftelse. EM was funded by the Deutsche Forschungsgemeinschaft (SFB 1218—269925409 and EXC 2030— 390661388). TP was supported by Knut och Alice Wallenbergs stiftelse and Vetenskapsra det (VR 2020-00884). MR was financially supported by the Knut och Alice Wallenbergs Stiftelse as part of the National Bioinformatics Infrastructure Sweden at SciLifeLab. PS was supported by the Stockholm City Council and Knut och Alice Wallenbergs Stiftelse.

Figures have been created using BioRender.com.

