## Supplementary figures and images for "Parkin is not required to sustain OXPHOS function in adult mammalian tissues"

### Fig. S1

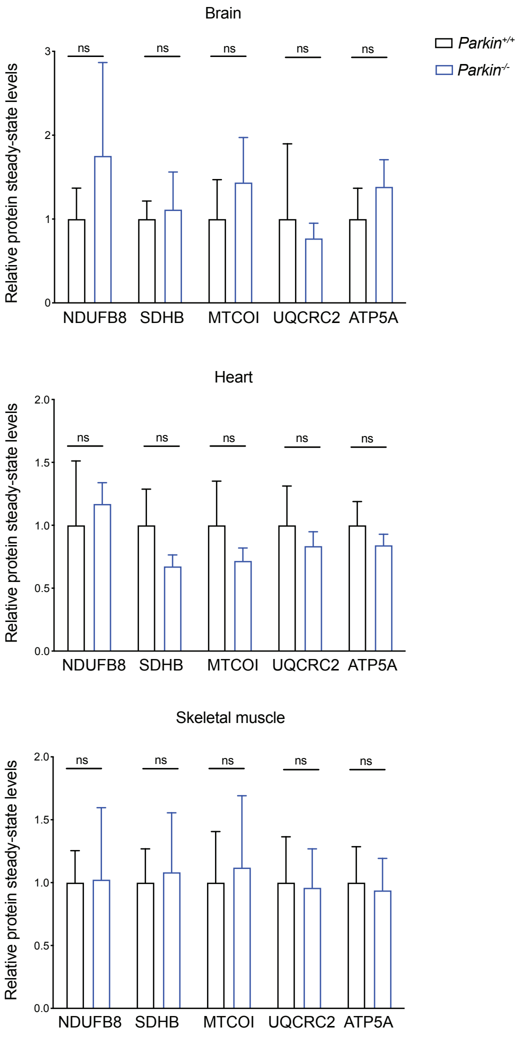

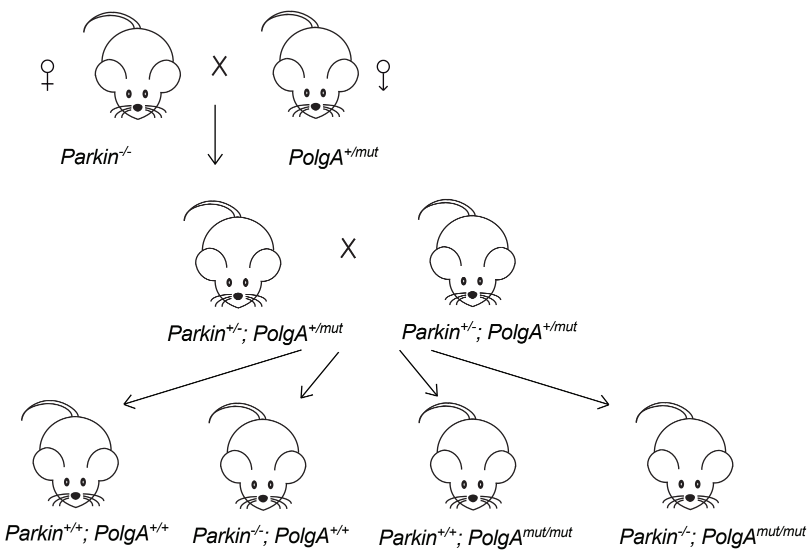

### Fig. S2

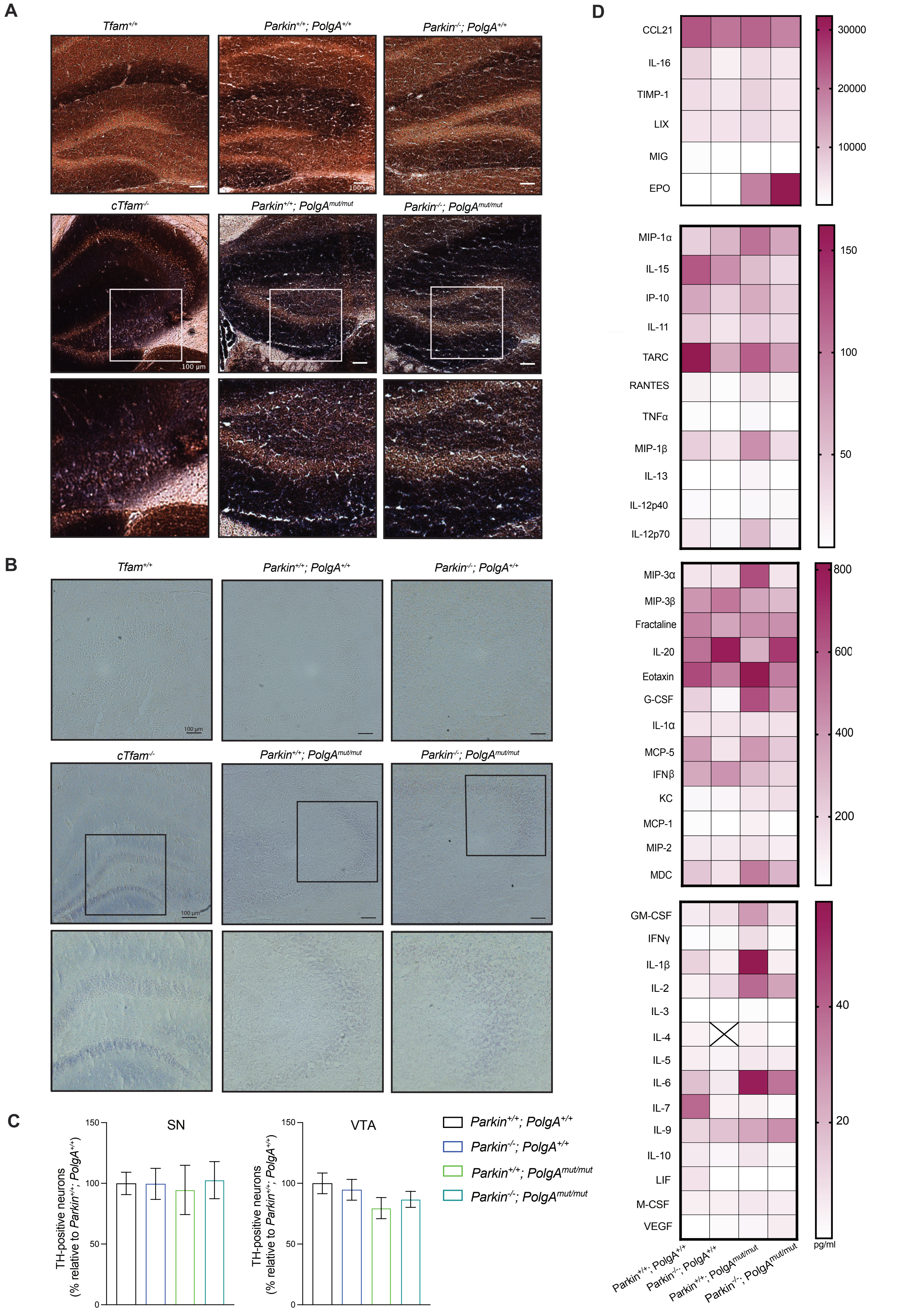

### Fig. S3

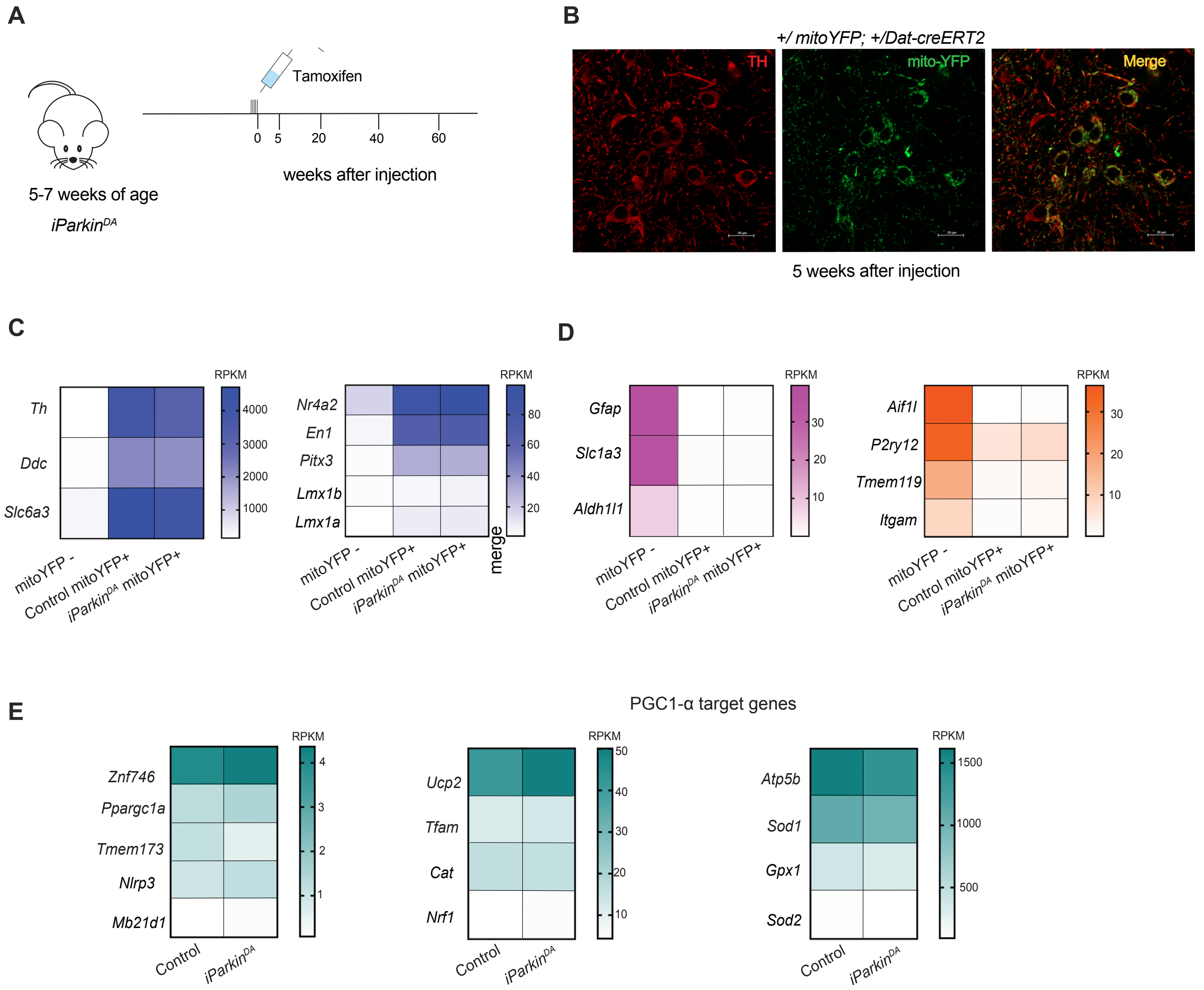
